# Mutational likeliness and entropy help to identify driver mutations and their functional role in cancer

**DOI:** 10.1101/354324

**Authors:** Giorgio Mattiuz, Salvatore Di Giorgio, Lorenzo Tofani, Antonio Frandi, Francesco Donati, Matteo Brilli, Silvestro G Conticello

## Abstract

Alterations in cancer genomes originate from mutational processes taking place throughout oncogenesis and cancer progression. We show that likeliness and entropy are two properties of somatic mutations crucial in cancer evolution, as cancer-driver mutations stand out, with respect to both of these properties, as being distinct from the bulk of passenger mutations. Our analysis can identify novel cancer driver genes and differentiate between gain and loss of function mutations.

---

Cancer genomes display a complex landscape determined by the accumulation of mutations. In individual cancer types specific patterns of mutations, referred to as mutational signatures^1^, have been identified, that can be ascribed to errors during DNA replication or repair, as well as to other effects of exposure to mutagens. However, in each tumour only a handful of mutations define the cancer phenotype and influence its evolutionary process (*driver* mutations)^2–4^. The majority of somatic mutations found in a given tumour are mostly neutral with respect to cancer evolution (*passenger* mutations), as they hitchhike on fitness-increasing mutations. In numbers, passenger mutations greatly exceed driver mutations, hence they can be used to describe the *neutral* mutational landscape of cancer genomes.

Identifying driver mutations in the haystack of passenger mutations is a major outstanding problem in cancer research. Several approaches to discriminate between driver and passenger mutations have been developed, based on factors such as mutation frequency^5–7^, gene expression^8^, protein domain analysis^9,10^, markers of positive selection^11^, network enrichment analysis^12^ and recurrent amino acid change analysis^13–16^.

Since driver mutations are under positive selection^11^, their mutational pattern might diverge from that observed in the much more numerous passenger mutations. In order to test this notion, we made use of a dataset of cancer mutations derived from the one generated by Chang et al.^16^ using several cancer-genome resources. The full dataset comprises ~2 million single-nucleotides variants present in over 11,000 cancer exomes from patients who had one of 41 tumour types. In order to calculate the probability of non-synonymous mutations, we applied on this dataset a Markov model trained on synonymous mutations, as they are mostly neutral. We preferred a zero-order rather than a higher order model, as we are dealing with coding sequences where, by virtue of the triplet genetic code, higher order patterns are confounded by constraints related to the protein sequence. Having worked out the parameters of the transition matrix of our model based on synonymous mutations, we refer to this output as the *Mutational Background Model* (see methods, **Fig.1a**), as these mutations reflect the outcomes of errors in the replicative/repair pathways and/or exposure to mutagens during cancer onset and progression. Next, we used the background model to calculate for each group of non-synonymous mutations (GNSM: the set of all mutations hitting the same codon in a given transcript among all patients) two scores. (a) *Mutational likeliness*: this measures the probability for a given GNSM to result simply from the background model. A negative value of this parameter indicates a nucleotide change that does not conform to the overall mutational pattern of the tumour; in other words, a decreased likeliness of an individual mutation may reflect selective pressure on that mutation. (b) *Mutational entropy*: this score, calculated by applying the Shannon entropy to each mutated codon, measures the bias towards a specific amino acid that is encoded by a GNSM compared to the expectations from the background model (**Fig.1a**). Mutational entropy is maximal when all amino acids are equally represented; entropy is zero when, of all possible outcomes, we observe only one amino acid.

**Fig.1.**
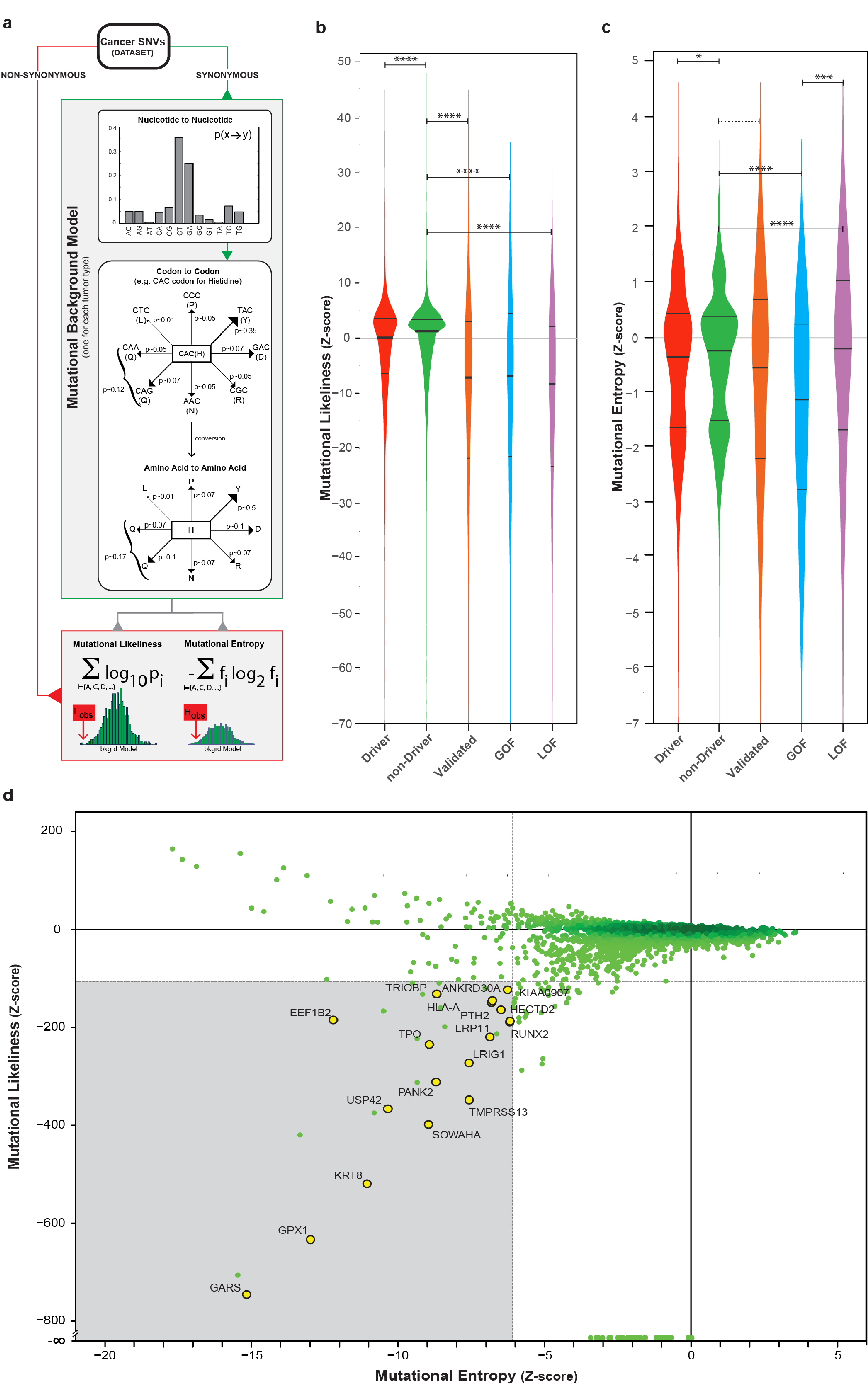
Mutational likeliness and mutational entropy distribution highlight differences between cancer driver and passenger mutations. (**a**) Synonymous mutations from the cancer dataset are used to derive the probability for each single nucleotide change in each tumour type (*Nucleotide to Nucleotide mode*l). The values obtained are used to build a model representing the probabilities for each non-synonymous change to occur given a tumour-specific mutational background (*Codon to Codon model*). From this, the probability for all the codons that can be produced by single nucleotide substitutions from any other codon was calculated (summing probabilities as needed; e.g. the case of Q) (*Amino Acid to Amino Acid model*). This model is then used to estimate the *mutational likeliness* and *entropy* scores for each non-synonymous mutation using the equations shown. p_i_ in the *mutational likeliness* formula represents the probability of a given amino acid change, depending on the wild type codon and the mutational background model. The *mutational entropy* is calculated considering the frequency (f_R_) of amino acid changes observed at the mutated codon. The *mutational likeliness* and *mutational entropy* were converted to Z-scores and the relative distributions were plotted. (**b, c**) The box-plots show the Z-score distributions of codons arising from cancer driver mutations (*Driver*, n=2666 positions on 399 genes), non-driver mutations (*non-Driver*, n=31037 positions on 9846 genes), experimentally validated ones (*Validated*, n=293), gain-of-function (*GOF*, n=100) and loss-of-function (*LOF*, n=172). and the horizontal black lines indicate the median for each group. The statistical significance is indicated *-p≤0.05; **-p≤0.01; ***-p≤0.001; ****-p≤0.0001; full details in the **Table 1**); the comparison not reaching statistical significance is indicated by a dotted horizontal line. (**d**) Scatterplot of mutational likeliness and entropy of the *non-Driver* dataset (**Supplementary Table S2**). Areas below the 1^st^ percentile of the *Driver* dataset (*mutational likeliness* = −113.68; *mutational entropy* = −6.16) are shaded in grey. Codon mutations on genes experimentally associated to cancer related processes are indicated in yellow.

**Table 1.** The p-values for each comparison are shown. Significant differences are reported in **Fig.1b** and **c.**

| Mutational Likelihood |  |  |  |
| --- | --- | --- | --- |
|  |  |  | Bonferroni (p-value) |
| *Driver* | vs | *non-Driver* | < .0001 |
| *Validated* | vs | *non-Driver* | < .0001 |
| *non-Driver* | vs | *GOF* | < .0001 |
| *non-Driver* | vs | *LOF* | < .0001 |
| *GOF* | vs | *LOF* | 1.000 |

| Mutational Entropy |  |  |  |
| --- | --- | --- | --- |
|  |  |  | Bonferroni (p-value) |
| *Driver* | vs | *non-Driver* | 0.0281 |
| *Validated* | vs | *non-Driver* | 0.0696 |
| *non-Driver* | vs | *GOF* | < .0001 |
| *non-Driver* | vs | *LOF* | < .0001 |
| *GOF* | vs | *LOF* | 0.0016 |

The significance of the two scores for each GNSM was then calculated by simulating 10,000 equally sized groups of mutations according to probabilities based on the background model. Then, for each GNSM we transformed mutational likeliness and mutational entropy into the corresponding Z-scores, using the estimated average and standard deviation obtained from the simulations (**Supplementary Table S1**).

In order to compile a list of non-synonymous mutations in *bona fide* cancer driver genes (*Driver*), and a list of non-synonymous mutations in other genes *(<u>n</u>on-Driver*: presumably passenger mutations) we used the Cancer Gene Census^17^. We use this information to compare the Z-score distributions of mutational likeliness and mutational entropy for the *Driver* and the *non-Driver* sets of mutations (**Supplementary Table S1**).

The distribution of mutational likeliness for *Driver* mutations is significantly shifted towards lower values than that of *non-Driver* mutations (p<0.0001, **Fig.1b**). This means that a substantial fraction of nucleotide changes in the driver genes are not explained by the mutational background model: we suggest this is due to positive selection acting on the cancer driver mutations. The shift is even greater (p<0.0001) when we only consider a set of validated driver codons (*Validated,* 293 positions on 75 genes, **Supplementary Table S1**), whose oncogenicity has been experimentally demonstrated (**Fig.1b**). This further reinforces the notion that the distribution of mutations that have been the subject of Darwinian selection during cancer development stand out as not representative of the overall outcomes of the mutational processes taking place in the cell during tumorigenesis or in the tumour.

With respect to mutational entropy, again the values we obtained are lower for *Drivers* than for *non-Drivers*, albeit with a lower level of statistical significance (p=0.0281, **Fig.1c**): this suggests that selection favours a reduced set of amino acid changes at these positions compared to the background model. However, we did not observe a significant difference between *Validated* and *non-Driver* mutations with regards to mutational entropy.

We reasoned that the entropy of a GNSM might be closely related to protein function: only a small subset of the amino acid changes that can be derived from a given codon will produce gain of function (GOF); conversely, a wider set of amino acid changes can lead to loss of function (LOF). For instance, among *TP53* mutations, A161T results in a gain of function (GOF)^18^, whereas R181P results in loss of function (LOF)^19^. Indeed, in our dataset, A161 mutations mostly change to Threonine, which results in an entropy score of −0.91. Conversely, there is no preference with regards to mutations at R181 and the entropy score is 0.08. When we analysed a whole set of *Validated* amino acid changes that are associated with GOF (n=100) or with LOF (n=172) (**Supplementary Table S1**), we found no difference between their mutational likeliness distributions (**Fig.1b**); on the other hand, the distribution of entropy values for GOF mutations when compared to that of LOF mutations was significantly shifted towards lower values (p=0.0016; **Fig.1c**). Thus, whereas the same evolutionary pressure applies to both GOF and LOF mutations (as they bear similar likeliness distributions), the difference in mutational entropy points towards the functional differences acquired through the amino acid changes. Thus, entropy scores highlight the dichotomy between gain- and loss-of-function mutations, which is fundamental in cancer biology, as in general GOF is characteristic of oncogenes and LOF of tumour suppressor genes.

Based on these results, we examined whether mutational likeliness and mutational entropy could provide a way to identify novel driver genes. We performed bibliographic searches on the 31 mutant genes in the *non-Driver* list for which mutational likeliness and entropy were below the 1^st^ percentile of the *Driver* distribution (**Fig.1d**; **Supplementary Table S2**): we found that 18 of them (58%) are convincingly associated with oncogenesis and/or cancer phenotypes (on 6 there is no information: see **Supplementary Table S2**). The same was true for the 68 out of 172 *non-Driver* genes within the 5^th^ percentile for which information was available (**Supplementary Table S2**). Among the 31 genes below the 1^st^ percentile, only one has been identified by other approaches^5,11,15–17^ (11 below the 5^th^ percentile) (**Supplementary Table S2**). Among these, one mutation in *RUNX2*^*20-21*^, that encodes a transcription factor associated with lymphomagenesis and bone metastasis, is not currently found in any of the cancer driver gene lists (**Supplementary Table S2**). Thus, our approach might be able to point towards a different set of cancer driving genes that may not be otherwise discoverable.

Cancer genomes are riddled with somatic mutations that are in part spontaneous and in part result from exogenous mutagens. Only a few of the mutant genes are subject to selection, and this has made it difficult to disentangle driver mutations from passenger mutations within the mutational landscape. We have found that likeliness and entropy of individual mutations can identify known driver mutations and predict some that have yet to be confirmed.

The difference in likeliness distributions between driver and passenger mutations is consistent with the notion that the mutational signatures observed in cancer genomes mainly reflect passenger mutations. Of course known cancer driver mutations may conform to the mutational background (e.g. *PIK3CA* mutations in HPV-related cancers^22^); but a low mutational likeliness score emerges as characteristic of cancer driver mutations, as they have been selected among the many others generated by the mutational processes prevailing in a particular tumour.

This analysis may also enable for the first time to differentiate between gain- and loss-of-function mutations. Whereas mutations are stochastic phenomena, and a set of mutations may bear the ‘signature’ of a mutagenic agent, evolutionary pressure depends on GOF, or LOF, or any change of function entailed by any particular mutation, regardless of its original signature.

Mutational likeliness and mutational entropy can identify cancer-driving mutant genes that are missed by other approaches and can guide the selection of potential cancer drivers for experimental validation. This is of special importance for precision medicine, since driver mutations are preferred potential targets of new therapies.

## Methods

### The cancer dataset

We started with a dataset of cancer mutations obtained from Chang et al.^16^, comprising of 2 million single-nucleotides variants (SNVs) identified in 11,115 cancer exomes from 41 tumours types. From the starting dataset, we removed the SNPs and all that SNVs that did not match the correct position in the coding sequence with regards to grch37 coding sequences. The dataset we used thus consisted of 1,799,208 mutations.

### Mutational background model

For all patients with more than 100 total mutations in coding regions, we selected the synonymous ones to calculate the transition matrix of a zero-order Markov model describing the background mutational process of each tumor type. All data from patients affected by the same tumor type contribute to the transition matrix for the given tumor type (Adrenocortical Carcinoma, Adenoid Cystic Carcinoma, Hypodiploid Acute Lymphoid Leukemia, Bladder Cancer, Breast Invasive Carcinoma, Cervical Squamous Cell Carcinoma And Endocervical, Chronic Lymphocytic Leukemia, Colorectal Carcinoma, Cutaneous Squamous Cell Carcinoma, Non Hodgkin Lymphoma, Esophageal Carcinoma, Gallbladder Carcinoma, Glioblastoma, High Grade Pontine Glioma, Head And Neck Squamous Cell Carcinoma, Kidney Chromophobe Cancer, Kidney Renal Clear Cell kirc Carcinoma, Kidney Renal Papillary Cell Carcinoma, Acute Myeloid Leukemia, Brain Lower Grade Glioma, Liver Hepatocellular Carcinoma, Lung Adenocarcinoma, Lung Squamous Cell Carcinoma, Lung Small Cell Carcinoma, Medulloblastoma, Mantle Cell Lymphoma, Myelodysplasia, Multiple Myeloma, Rhabdoid Cancer, Neuroblastoma, Nasopharyngeal Carcinoma, Adenocarcinoma Ovarian Serous Cystadenocarcinoma, Pancreatic Adenocarcinoma, Pancreatic Neuroendocrine Carcinoma, Pilocytic Astrocytoma, Skin Cutaneous Melanoma, Stomach Adenocarcinoma, Thyroid Carcinoma, Uterine Corpus Endometrial Carcinoma, Uterine Carcinosarcoma, Prostate Adenocarcinoma). For cancer types for which the transition model could not be calculated as there were no tumours with at least 100 mutations we built an average model obtained by averaging all models from the other cancer types.

As expected, considering the directionality of coding sequences, complementary mutations (e.g. A → C and T → G) are not symmetric and we treated them separately. Each model is thus composed by a vector of 12 probabilities, one for each possible nucleotide change:

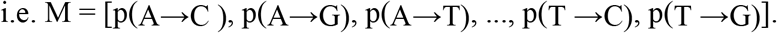

As we analyzed mutations in their codon context, we define as a group of non-synonymous mutations (GNSM) the set of all mutations hitting the same codon in the same transcript in different patients. We considered for analysis only those codons for which at least three mutations existed in the dataset.

### Mutational likeliness

The mutational background model is used to extrapolate the background probability distribution of amino acids resulting from non-synonymous mutations hitting a certain codon (**Fig.1a**).

This distribution is then compared with the set of amino acids observed at a given codon mutated. For instance, let’s suppose that several patients have a given codon CAC (Histidine) mutated. From the background model we know the values of p(C→A, G, T) and p(A→C, G, T) and therefore we can calculate the probability of going from one codon to another by means of a single point mutation, e.g. p(CAC → AAC, GAC,…). From the probabilities towards each possible codon, we calculate the corresponding expected distribution of amino acids by merging the probability of codons coding for the same amino acid. In the case of the CAC codon we get all the possible resulting codons, which lead to N, D, Y, P, R, L, Q, Q. Thus, each amino acid change has its own probability to happen in this codon context:

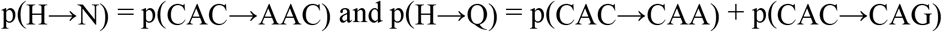

Since we only consider non-synonymous mutations, while some of the amino acids reachable from a certain codon are synonymous, the probabilities for non-synonymous changes are then rescaled to 1.

We define the mutational likeliness score as:

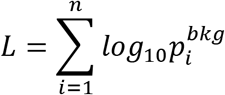

with *n* the number of observed mutations and 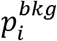 the probability of a certain mutation given by the background model; this formula therefore allows to calculate the probability of a certain set of mutations at a certain codon for a specific tumour background model. Mutations in line with the background model will have large *mutational likeliness,* as they will tend to have large 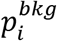, while those that do not conform to the background model show a bias towards smaller scores. We then perform random sampling to assess the significance of the score observed for each GNSM: we use the background model to generate 10,000 equally sized sets of mutations starting from the wild type codon and we calculate the average and standard deviation of the score. This procedure allows to calculate the Z-score and therefore the significance for each observed GNSM. This measure allows the identification of the codons where the set of observed mutations diverges the most with respect to the background model.

### Mutational entropy

In order to analyze whether (a subset of) driver mutations tend to prefer certain amino acid changes, we consider all amino acids changes found in each patient genome for a given GNSM, and we calculate its entropy using the Shannon formula^23^:

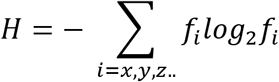

Where the sum runs over the different amino acids encoded by the GNSM (x, y, z…) with frequencies *f*_*i*_. As in the previous case, the entropy of each set of mutations is transformed in a Z-score by using the background model to produce 10,000 equally sized group of amino acids from which we calculate the expected average entropy and its standard deviation. Therefore, while the *mutational likeliness* identifies mutations with a pattern of nucleotide changes differing significantly from the expected (given the background model), the *mutational entropy* identifies those mutations that might be in line with the background probabilities but not with the expected amino acid distribution.

### Cancer Gene Census List

We exploited the Cancer Gene Census^17^ to define a list of known cancer driver genes. In this work, we excluded all those genes that are not present in our dataset and whose oncogenicity derives from copy number alterations, gene fusions and truncations, insertions, or deletions. Our *Driver* dataset comprises of 2666 mutations on 399 driver genes (**Supplementary Table S1**). The complementary dataset of mutations outside these driver genes (*non-Driver* includes 31037 mutations on 9846 *non-Driver* genes (**Supplementary Table S1**).

### Experimentally Validated Driver Codons (*Validated*)

Starting from our list of 2666 driver mutations, we manually selected 293 codons for which the effects of the specific mutation on the gene have been reported in the literature. Many of the validated positions were taken from the JAX Clinical Knowledgebase (The Jackson Laboratory; https://ckb.jax.org) and confirmed through further bibliographic search. We divided the *Validated* amino acid changes in two subgroups containing mutations inducing either gain-of-function (GOF = 100) or loss-of-function (LOF = 172) depending on the effect of the amino acid change on protein function. To create GOF and LOF lists, we usually had to consider only the starting amino acid position (e.g. BRAF V600); in rare cases a single codon could be classified both as GOF or LOF, depending on the resulting amino acid change; in our analysis we considered each mutation separately, in both subgroups (e.g. the mutation of Y646 in the EZH2 gene leads to LOF for Y646C or to GOF for Y646F) (**Supplementary Table S1**). It is noteworthy that only 10% of mutations in the *Drivers* dataset has been tested and catalogued as GOF or LOF. This highlights the need for experimental approaches to validate cancer-associated mutations and to allow a proper identification and functional characterization.

### Driver mutations in the *non-Driver* dataset

We selected mutations in the *non-Driver* dataset whose *mutational likeliness* and *entropy* scores were below the 1^st^ and the 5^th^ percentiles of the *Driver* dataset distributions. For each gene, we performed a bibliographic analysis (using all aliases for the gene name) and selected those whose function had been linked to cancer processes and phenotypes. (**Supplementary Table S2**).

### Statistical analysis

The distributions of the mutated codon groups were compared using the Mann-Whitney test with Bonferroni multiple comparison correction. Continuous variables were expressed as mean, standard deviation, median, 25^th^ and 75^th^ percentiles. The significance level was set to 5%. The statistical analysis was performed using SAS 9.3.

## Authors Contribution

GM, AF, MB, and SGC conceived the study; GM, AF, MB, and SGC wrote the manuscript; MB developed the mathematical model; GM, SDG, and MB performed the bioinformatics analysis; LF performed the statistical analysis; GM, FD performed the bibliographic analysis.

## Acknowledgements

We thank Rosario Notaro for the intelligent discussions that nudged us into this work. We are deeply indebted with Lucio Luzzatto, as his many rounds of critical reading and commenting improved the focus and the clarity of the text. This work was supported by a grant from the Italian Ministry of Health (PE-2013-02357669); GM and FD were supported by a grant from AIRC (IG-17701).

